# Collective detection based on visual information in animal groups

**DOI:** 10.1101/2021.02.18.431380

**Authors:** Jacob D. Davidson, Matthew M. G. Sosna, Colin R. Twomey, Vivek H. Sridhar, Simon P. Leblanc, Iain D. Couzin

## Abstract

The spatio-temporal distribution of individuals within a group (i.e its internal structure) plays a defining role in how individuals interact with their environment, make decisions, and transmit information via social interactions. Group-living organisms across taxa, including many species of fish, birds, ungulates, and insects, use vision as the predominant modality to coordinate their collective behavior. Despite this importance, there have been few quantitative studies examining visual detection capabilities of individuals within groups. We investigate key principles underlying individual, and collective, visual detection of stimuli (which could include cryptic predators, potential food items, etc.) and how this relates to the internal structure of groups. While the individual and collective detection principles are generally applicable, we employ a model experimental system of schooling golden shiner fish (*Notemigonus crysoleucas*) to relate theory directly to empirical data, using computational reconstruction of the visual fields of all individuals to do so. Our integrative approach allows us to reveal how the external visual information available to each group member depends on the number of individuals in the group, the position within the group, and the location of the external visually-detectable stimulus. We find that in small groups, individuals have detection capability in nearly all directions, while in large groups, occlusion by neighbors causes detection capability to vary with position within the group. We then formulate a simple, and generally applicable, model that captures how visual detection properties emerge due to geometric scaling of the space occupied by the group and occlusion caused by neighbors. We employ these insights to discuss principles that extend beyond our specific system, such as how collective detection depends on individual body shape, and the size and structure of the group.

## 1 Introduction

Being part of a group is an effective strategy for avoiding predation threats [1–4] and locating promising resources [5, 6]. Enhanced detection of external objects (for example a predator, or a source of food) is a key aspect of being part of a group, with the benefits referred to as the ‘many eyes’ effect [7, 8]. The structure within a group influences how individuals interact with one another and the surrounding environment. For example, groups tend to have more individuals and an increased density under heightened predation risk [9–15] (but see [16, 17]). An individual’s position within the group can determine both its possible risk to predation [18], as well as the extent of its social interactions [19, 20]. Despite the importance of social grouping for gathering information about the external environment [21, 22], there has been little quantification of how within-group structure and the size of the group influence the group’s interactions with their environment.

Many species who form coordinated, mobile groups employ vision as a primary modality for mediating social interactions [23–25]. Visual connectivity among individuals can predict how a social contagion spreads through a group, such as when ‘informed’ individuals detect and move towards a cue associated with food, and are followed by other naive group members [19, 26], or when a startle response propagates across a group [14, 20]. As groups get larger, occlusion due to neighbors means that individuals differ in the visual information they have available to them. The available visual information determines whether individuals will respond to other group members [19, 20], as well as if the any individuals in the group will have the ability to detect cryptic stimuli, such as a predator [7, 8].

Here we examine how the visual information available to individuals in a group depends on both the number of group members and on how individuals are positioned within the group (i.e. the group’s internal structure). We first analyze, quantitatively via computational visual field reconstruction [14, 19, 20], the visual information available to all individuals within groups of golden shiner fish, whose social behavior is predominantly mediated by vision [20]. We performed experiments with groups of different numbers of fish, ranging from 10 to 151 in number. We examine how the detection coverage, which is the angular fraction of the external visual area that an individual can see, depends on the number of group members and an individual’s position within the group. We then formulate a simple model of external detection ability which demonstrates a good match to the main features observed in the data. The model generalizes to show how detection scales when a group contains more individuals and we use these results to discuss the implications and generalizations to other animal groups.

## 2 Results

We filmed free-schooling groups of 10, 30, 70, and 151 golden shiner fish (*Notemigonus crysoleucas*) in the laboratory and used a combination of automated and manual tracking to extract positions and orientations while maintaining individual identities over the course of each trial (see Methods). Golden shiners are a widespread species of freshwater fish [27] that are surface feeders and thus swim close to the surface of the water [28]. We estimated the external visual detection capabilities of each individual using a procedure where a neighboring individual can block the external vision of a focal individual in a certain direction (Fig 1A-B; see Methods). Individuals tend to have a ‘blind angle’ to the rear, which for this species has been determined to be 25 degrees [29], and we include this in the visual detection procedure (Fig 1C). In addition, we note while individuals form a relatively planar group structure, near the surface, the arrangement is not perfectly 2-dimensional; this means that neighboring individuals do not always block an external view. Since our tracking is only in 2D, we account for out-of-plane effects by considering incomplete visual blockage, where each neighbor has a certain probability of blocking external vision (Fig 1D). Applying the detection algorithm to each individual in the school illustrates the overall external detection abilities of the group (Fig 1E-F; [30]).

**Figure 1:**
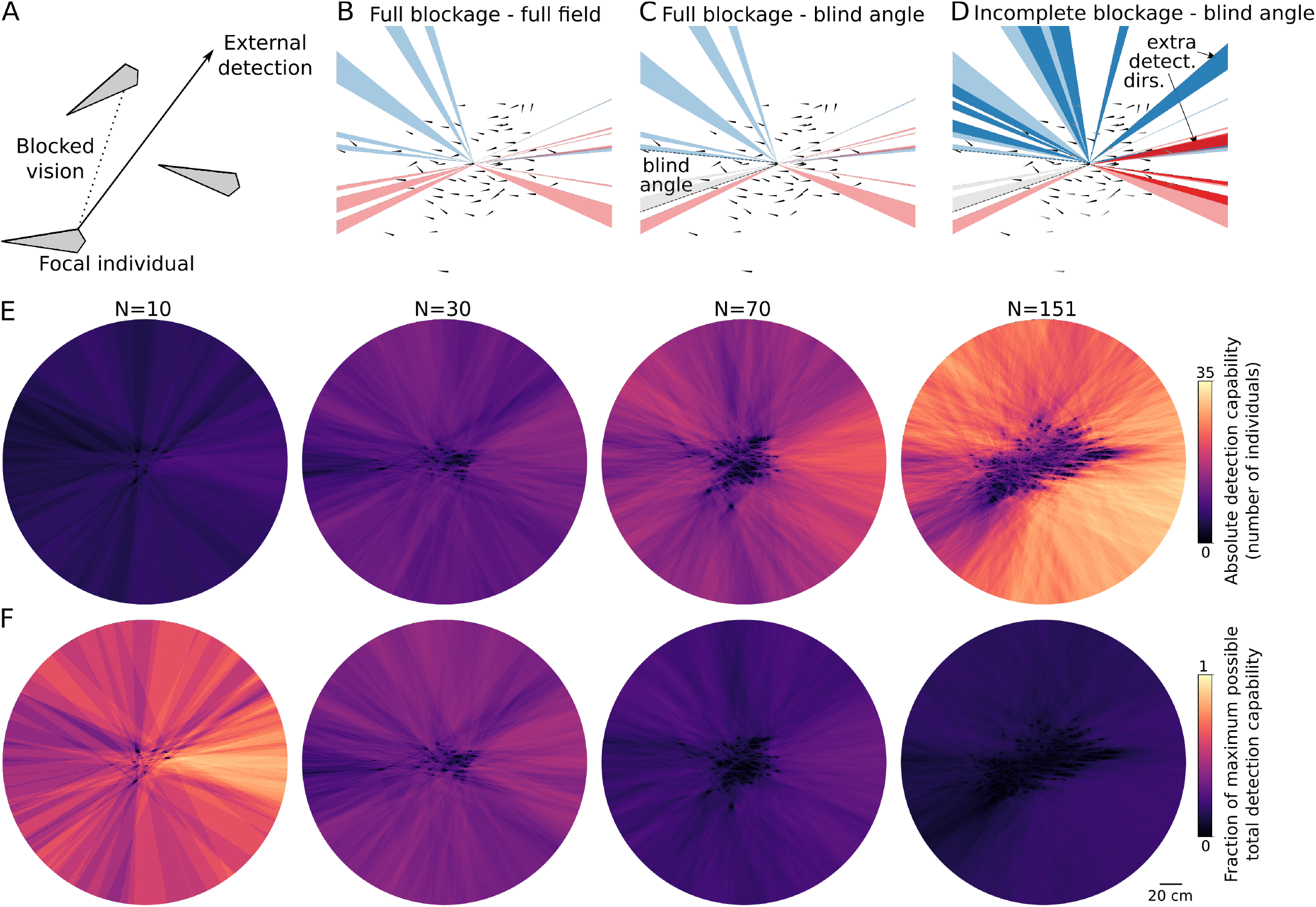
External visual detection. (A) A focal individual has external detection in a given direction if neighbors do not block vision in that direction. (B-D) External visual detection coverage of a single individual located at the center of a school of 70 fish. Shown is the detection coverage determined using different parameters. Directions where the left eye has external detection capability are shown in blue, and that for the right eye in red. (B) Full visual blockage and a full 360° field of view. (C) Including a blind angle where fish cannot see behind themselves, with otherwise full blockage from neighbors. The blind angle area is highlighted by the dotted lines, and detection directions omitted due to the blind angle are shown in gray. (D) Blind angle and incomplete (probabilistic) blockage, using a blockage probability of 75%, where some neighbors do not block the external view in a certain direction. Neighbors that are ignored are shown in gray, and additional detection directions are shown in darker colors. (E-F) Illustration of the external visual field of the entire group at a single frame. The heatmap shows detection capability obtained by summing the overlapping regions of the external visual fields of all individuals, using results with incomplete blockage (75%)-blind angle. Results are displayed by scaling to show either (E) Absolute detection capability in terms of the number of individuals with detection capability, or (F) The fraction of the maximum possible total detection capability among group members.

### 2.1 Individual detection coverage

We first examine individual detection coverage, which ranges from 0 to 1 and represents the fraction of the external visual space that an individual can see, and then following this, in Section 2.2, examine the total number of group members with detection capability in a certain direction at a moment in time. For small groups of 10, all individuals have a large detection coverage and can see nearly the full range around the group, i.e. in directions to the front, back, and side of the group, regardless of their position within the group. As the number in the group increases, however, the average detection coverage decreases due to occlusion caused by neighbors. Additionally, the variance of individual external visual coverage in the group increases with the number of individuals, reflecting an increased heterogeneity in visual access resulting from individuals of the group having their visual field increasingly dominated by others, thus occluding their view of areas external to the group (Fig 2). Considering a blind angle decreases the instantaneous detection coverage, with the largest effect for the group of 10. This is because in in small groups, the rearward area, in the absence of a blind angle, would be visible, while in large groups, it is likely that vision to the rear is already blocked by a neighbor.

**Figure 2:**
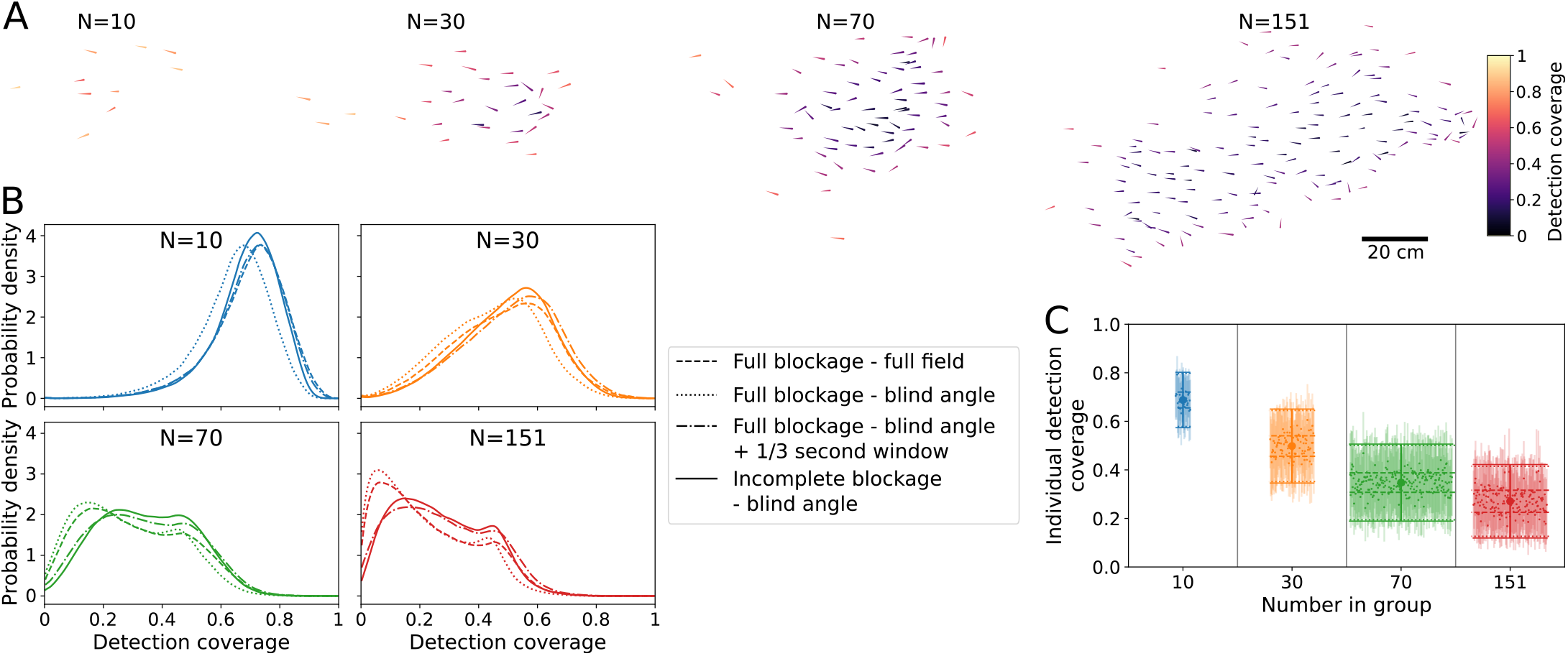
Individual detection coverage. The detection coverage is the fraction of the external visual field that an individual can see. (A) Example snapshot of the external detection coverage for groups with different numbers of individuals. (B) Distributions of individual detection coverage for the different groups, combining all individuals during a trial, calculated using different settings: full blockage-full field (dashed line), full blockage-blind angle (dotted line), full blockage-blind angle with detection capability any time over a 1/3 second time window (dashed-dotted line), and incomplete blockage (75%)-blind angle. (C) Detection coverage, comparing individual differences to the combined distribution. Results use incomplete blockage (75%)-blind angle. The mean and standard deviation of the combined distribution from B is shown as the large point and error bars for each group size. The mean individual detection coverages during a trial are the small points, and individual standard deviations are the shaded bars. Note that 3 trials were performed for *N* = (10, 30, 70), while only 1 trial was performed for *N* = 151. The individual points are spaced on the x-axis for display purposes. The dashed line shows the contribution from “consistent individual differences” (the variance of individual means) to the overall distribution, while the dotted line shows the contribution of “individuals differing during a trial” (the mean of individual variances; see Methods, Eq. 7).

In addition to instantaneous coverage, we consider that since individuals move over time, small positional changes may increase the effective range of visual information that is available during a certain time window. To represent this, we say an individual has detection capability in a certain direction at time *t* if there was visual access in that direction at any time within the previous *T* seconds, i.e. within the time window of *t* − *T* to *t*. Using a time window increases the average detection coverage, with the largest effect on the most numerous group (*N* = 151). For all groups, the results using a blind angle and a window time of *T* = 1/3 sec. yield average detection coverage values that are near to or greater than that without using a blind angle. This demonstrates that considering small positional changes over a short time can effectively “mitigate” the decrease in detection coverage caused by having a blind angle.

As expected, considering incomplete blockage at an instant in time increases an individual’s detection coverage, with the largest shift for the most numerous group (*N* = 151). Fig 2 shows results with a blockage probability of 75%; using other values causes the coverage to progressively increase as the probability of individuals blocking one another decreases. Despite the shifts in distributions when considering a blind angle and incomplete blockage, we see the same the main trends: individual detection coverage decreases and the variance of the distribution of individual detection coverage increases when there are more individuals in the group (Fig 2). Because we know that out-of-plane effects lead to incomplete blockage, and that individuals do have a blind angle, in the following we focus in detail on the instantaneous detection results using a blind angle and incomplete blockage. Although we don’t have an exact value of what the effective blockage probability due to non-planar effects would be, we use the intermediate value of 75% blocking probability as a reasonable value and proceed by focusing on this case, noting that none of the general features of the results are dependent on the exact value of the blocking probability.

The distributions in Fig 2B combine all individuals over each trial. Are there consistent differences in detection coverage among particular individuals during a trial? Fig 2C shows that while both individual differences and changes during a trial contribute to detection coverage, consistent individual differences explain a much smaller fraction of the overall variance, in comparison to individuals changing their position in the group during a trial. Consistent individual differences explain on average only 8% of the total variance.

### 2.2 Group detection capability

Instead of examining detection coverage of individuals, we can instead ask about the total number of group members with external detection capability in a certain direction at a moment in time. This can depend on the group state and group area, e.g. whether the group is swimming in a polarized, milling, swarm, or other configuration (Fig 3A; [31]), as well as on the external direction with respect to the group travel direction. We first examine the former. The total number of possible external detections among all group members in any direction increases with *N*, and only shows a small dependence on the group state (Fig 3B). While the polarized state is the most common configuration - the groups of 10 and 30 do not spend significant time in the milling or swarm states - the groups of 70 and 151 do spend time in different states, and display slightly higher external detection abilities in the polarized state compared to milling or swarm states (Fig 3B).

**Figure 3:**
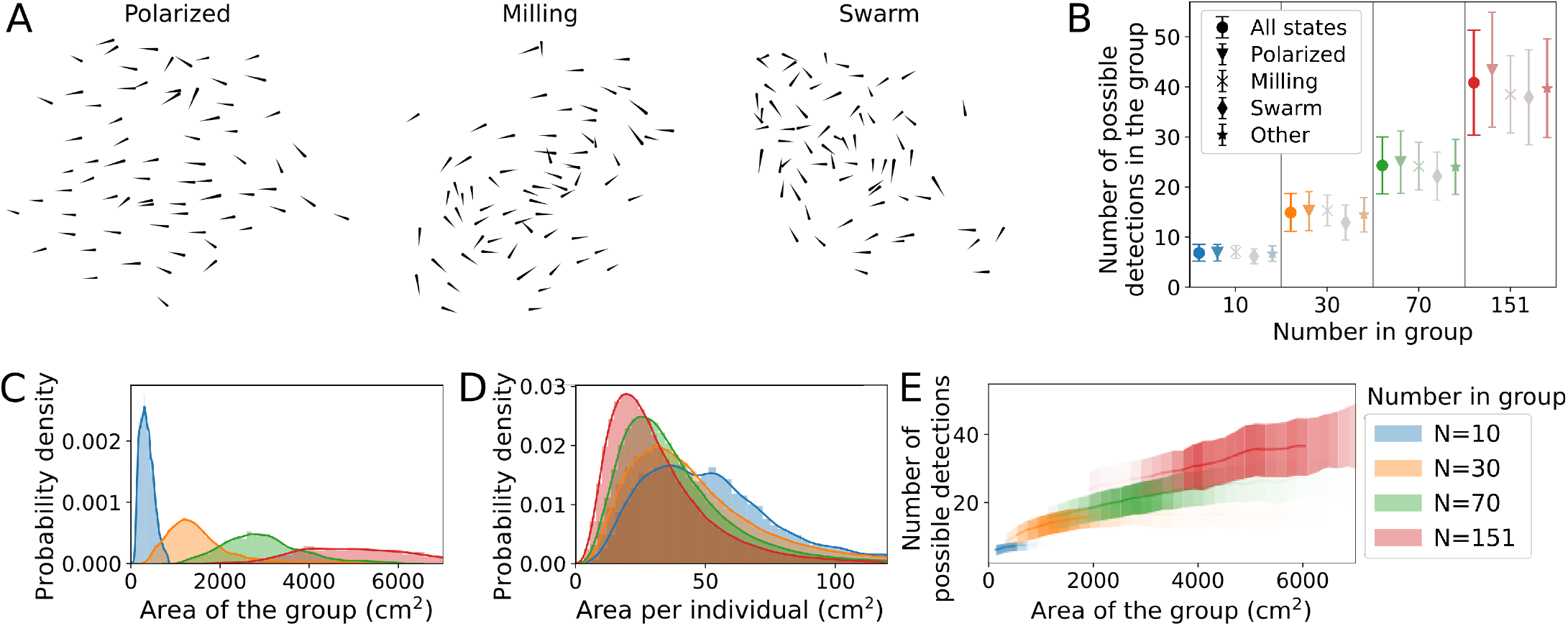
Group detection and state dependence. (A) Example snapshots of a group of 70 in different configurations: polarized, milling, and swarm states. (B) The total number of possible detections for all individuals in the group, for groups with different values of *N*, showing results for all group states compared to polarized, milling, swarm, and other states. Error bars show the standard deviation of the total number of possible detections among all group members in a certain direction at an instant in time (Eq. 9). The color saturation of the points is proportional to the amount of time a group of size *N* spends in a certain state, i.e. states not used much are shown in gray. Groups of *N* = 10 and *N* = 30 do not spend significant time in the milling or swarm states, and therefore these points shown in gray. (C) Distributions of the total spatial area occupied by groups of different numbers of individuals. (D) The spatial area per individual, calculated using Voronoi tesselation, for groups of different numbers of individuals. See Fig 6 for information on how the total group area and the individual area are calculated. (E) The total instantaneous detection capability among all group members, averaged over all possible directions over time, plotted as a function of the total area of the group at that time. The line shows the mean and the shading shows the standard deviation of the number of possible detections. The transparency of the lines is proportional to the probability that the group has a certain area value (see distributions in C).

In addition to swimming in different configurations, the density of the group can differ, for example with a dense vs. tightly-packed group configuration. Naturally, groups with more individuals occupy a larger spatial area. The standard deviation of the spatial area occupied is also larger for groups with more individuals (Fig 3C). The average spatial area per individual slightly decreases for larger *N*, reflecting that although the distributions were overlapping, individuals tend to pack slightly more tightly when more individuals are in the group (Fig 3D). The number of possible external detections among group members is higher when the group occupies a larger area; this is because when individuals are spaced further apart, each neighbor subtends a smaller angle on the visual field of others and therefore blocks less of the external view (Fig 3E).

### 2.3 Angular dependence of detection

The number of individuals with detection capability in a certain direction also depends on the angle with respect to the group travel direction. Note that if the group is not moving, then there is no ‘group travel direction’, and no front or back of the group. However, if the group is moving cohesively in a polarized configuration (see e.g. Fig 3A), then there is a clear travel direction and a difference between individuals at the front versus the rear of the group. Because of this, we consider only movement when in a polarized state to examine the angular dependence of detection [31]. For fish, which like many animals have elongated body shapes, detection capabilities are higher to the front of the group than to the side of the group. Due to the blind angle, detection capabilities are lowest to the rear of the group (Fig 4A; [30]).

**Figure 4:**
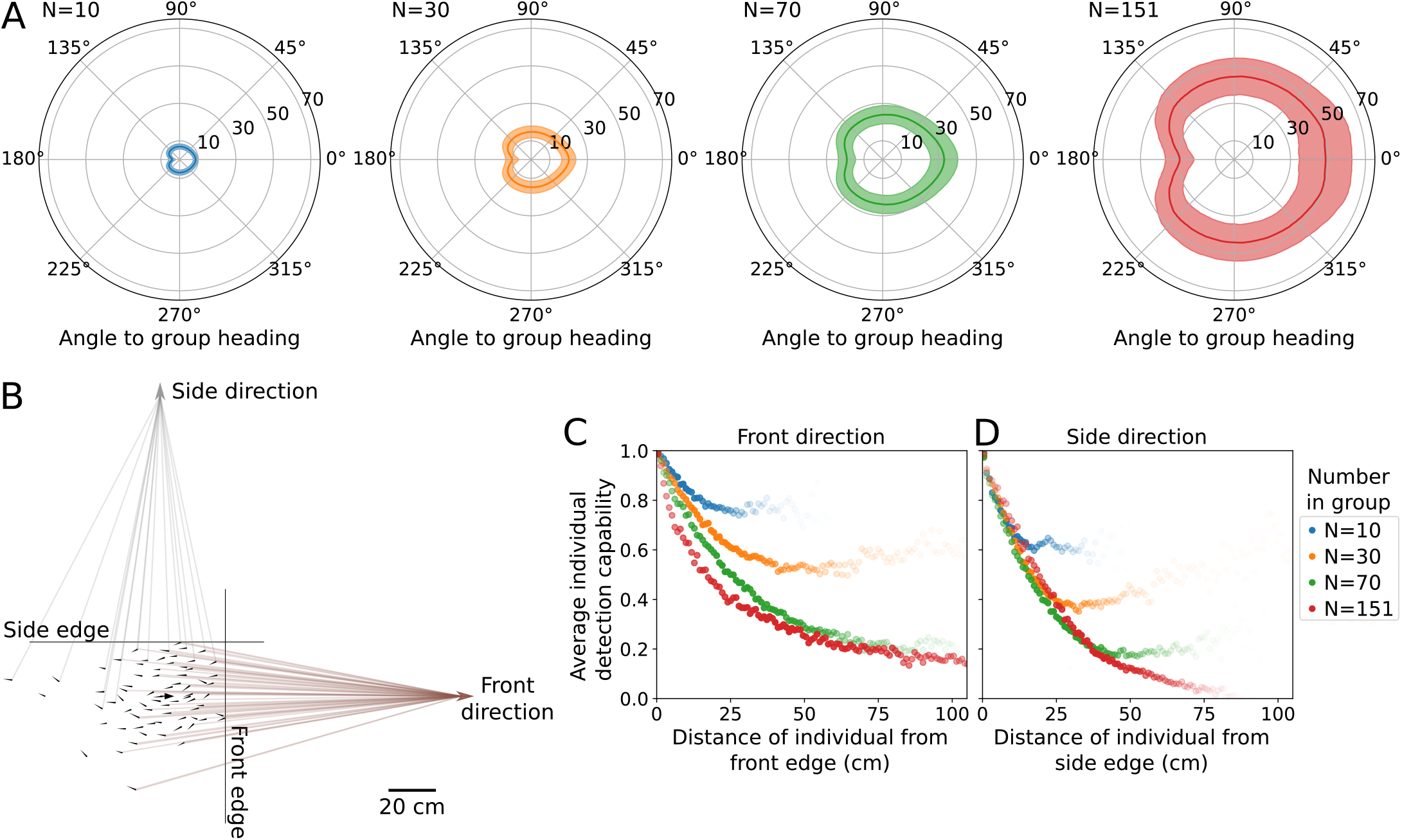
Detection relative to group heading direction. (A) The number of possible detections in the group at different angles with respect to the group heading direction. Results are calculated using instances when the group is moving aligned in a polarized state. (B) Illustration of front and side detection using a snapshot of a group of 70, showing where individuals have open lines of sight to either a location to the front (brown lines) or to the side (gray lines) of the group. The front and side edges of the group are defined by the furthest individual in these respective directions. (C) Average individual detection capability in a direction to the front of the group, plotted as a function of individual distance from the front edge of the group. (D) Analogous results to (C), but for average individual detection capability in a direction to the side of the group, plotted as a function of individual distance from the corresponding side edge of the group. For (C,D), the transparency of the points is proportional to the number of observations of individuals at that distance from the front or side.

We next examine how detection capability to the front or the side of the group depends on an individual’s in-group position. In-group position is represented by defining the ‘edge’ of the group as the individual furthest away from the centroid in that direction, and then calculating an individual’s distance from either the front or side edge of the group (Fig 4B). An individual located at a certain edge always has detection capability in the corresponding direction, and therefore average detection capability is 1 at a distance of zero from the edge. Detection capability then decreases with distance from the edge. However, the decrease in detection capability with distance from the edge depends on both the angle with respect to the group travel direction and the number of individuals in the group. In the smallest group (*N* = 10), nearly all individuals have detection capabilities to the front of the group, and the detection capability shows only a small decrease with distance from the front edge. The steepness of decay of detection capability with distance from the front edge of the group increases with the number of individuals in the group (Fig 4C).

In contrast, the detection capability with respect to the distance from the side edge of the group shows a similar initial decay for all group sizes, but extends ‘further’ for the larger groups because they take up a larger area (Fig 4D). This difference between front versus side detection is due to the elongated body shapes of fish as well as the alignment of individuals when swimming as a polarized group. While an individual’s vision to a region to the side of the group may be completely blocked by a single nearby aligned neighbor, visual blockage to a region to the front is more likely to depend on the positions and orientations of several neighbors [30].

### 2.4 Model of external detection

To better understand how detection capability changes with the number of individuals in the group, and to seek general principles, we formulate a simple model of external visual detection capability of a group of individuals. In the model, a group of *N* individuals occupies a circular area with radius *R*, within which there is a constant visual blockage probability of λ. At a distance *r* from the center, the probability of having detection capability at an angle of *θ* is proportional to the blockage probability multiplied by *g*(*r, θ*), which is the distance to the edge of the group in that direction (Fig 5A; see Methods). For a group with *N* individuals, we specify that the visual blocking probability scales according to

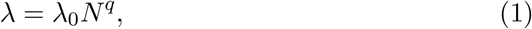

where λ_0_ is a baseline blocking probability and *q* is a scaling exponent. We fit the values of λ_0_ and *q* by comparing individual mean detection probabilities from the model to the data (Fig 5B).

**Figure 5:**
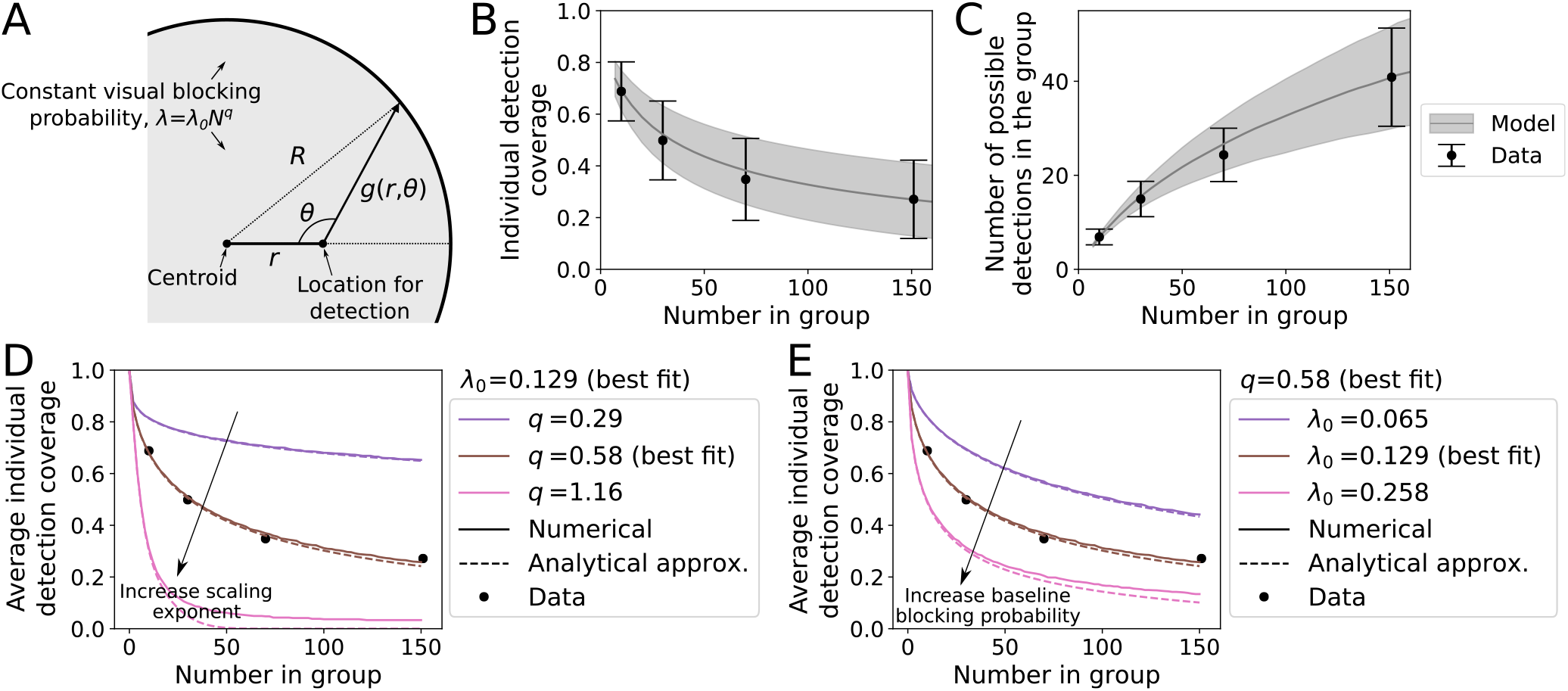
Model of external visual detection coverage. (A) Illustration of the quantities in the model. (B) Individual detection coverage in the model compared to the data. The baseline blockage probability λ_0_ and the scaling exponent *q* are fit to the average detection coverage in the data, yielding λ_0_ = 0.129 and = 0.58. The error bars show the standard deviations of the distributions from the data, and the gray shaded area shows the standard deviation for the model. (C) Total number of group detections in the model compared to the data. The parameter *σ*, which represents the standard deviation of the radius of the group in the model, is fit to the standard deviation of the number of possible detections in the data (error bars), leading to *σ* = 0.263. The gray shaded area shows the standard deviation of group detection in the model. (D-E) Average detection capability for different model parameters and number of individuals in a group. In each, the points show the values from the data, the solid lines are obtained numerically from the model, and the dashed lines are the series approximation in Eq. 2. The solid brown line shows the best fit from model, which is obtained using numerical evaluation. (D) Average detection capability for different values of the scaling exponent *q*, with λ_0_ set to the best fit value. (E) Average detection capability for different values of the baseline blocking probability λ_0_, with *q* set to the best fit value. See Methods for model details and fit procedure.

We furthermore include the effect that a group may change the spatial area it occupies by using a parameter *σ* for the standard deviation of the radius of group. Since a given group occupying a larger area has a higher average number of possible detections (Fig 3E), spatial area changes increase the standard deviation of the total number of possible detections of group members in a given configuration. We therefore fit the parameter *σ* to the standard deviation of the number of possible detections for the different group sizes (Fig 5C).

An approximate solution for the average detection coverage is obtained using a series expansion (see Methods), yielding

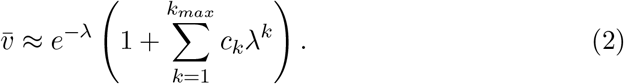

where 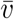 is the average detection coverage, λ is specified in Eq. 1, and the numerical values of the coefficients *c_i_* are listed in Methods from an expansion to 6^th^ order terms. Eq. 2 shows that to leading order, average detection decays exponentially in λ. From this, we clearly see that increases in both the scaling exponent (*q*) or the baseline blocking probability (λ_0_) both decrease the average detection capability. However, these two parameters affect the shape of the decrease differently, with *q* having more of an effect on the shape of the exponential decay with increasing *N* (Fig 5D-E).

By using different parameters, the model can generalize to describe the detection coverage of individuals at different densities, of different sizes, or with different scaling properties with *N*. A higher value of the baseline blocking probability λ_0_ could represent larger individuals, or a higher average density for a given value of *N*. The parameter *σ* represents the standard deviation of the spatial area that the group occupies about the average; since density affects detection coverage, a higher value of *σ* means a higher variance in the number of possible group detections. The parameter *q* represents how individual detection coverage changes with the number of individuals in the group (*N*); in particular, a higher value of *q* could represent groups that show a sharper decrease in the area per individual with *N* than we observed experimentally.

Overall, by assuming a simple shape of the group and a constant blockage probability, the model demonstrates that our experimental findings for detection results can be explained by the geometry of how neighboring individuals occlude an external view. In particular, the model can replicate the experimental observations that average individual detection decreases with *N* (due to an increased probability of occlusion from neighbors), and that the variance of the number of possible detections in the group increases with *N* (due to the variance in the spatial area occupied by the group).

## 3 Discussion

In general, in small groups individuals have detection capability in nearly any direction, while in large groups individuals can differ substantially from one another in their visual information due to occlusion from neighbors. For our study system (golden shiner fish), we thus find that meaningful distinctions in available visual information emerge when groups contain between 30-70 fish; at these sizes and larger, some individuals may detect an object while others do not. While previous work has used position-based metrics to define, for example, “peripheral” versus “central” locations within a group (e.g. [18, 22, 32]), we note that these distinctions are only meaningful with respect to detection for groups with sufficiently many individuals.

The density of a group affects visual detection abilities (Fig 3E). What functional aspects may lead a group to adopt a certain density? Previous work has suggested that up to a certain point, higher group density is associated with an increased ability to spread information among group members; or in other words, denser groups tend to be more ‘tightly connected’ [19, 20]. The reason for this, as has been found for animals from fish [20] to humans [33–35], is that the spreading of social behavior is best described by a fractional contagion process, whereby an individual’s probability of response depends on the fraction of their neighbors that have responded [36]. In denser groups, each individual has fewer neighbors —and thus a contagion is more likely to spread [19, 20]. This is further supported by what happens when fish are exposed to Schreckstoff, a chemical that is typically released to signal that a predator is nearby. In this case, individuals move closer together, which facilitates an increase in their ability to spread behavioral change, and thus exhibit a greater responsiveness to external threats [14]. While transmission of behavioral change among group members may be enhanced at higher density, our results show that external detection is enhanced at lower density (Fig 3E). This is because at lower density, neighbors subtend a smaller angle in the visual field of others. In the model, an overall lowering of the density of the group can be represented by a lower value of λ_0_. The ‘trade-off’ between external detection and internal communication may be a driver of the optimal group density, and can explain why the overall spatial area of the group does not predict how quickly a group will respond [37]. At an individual level, a low external detection capability to the side of the group tends to be compensated by stronger visual connectivity to neighbors [30].

With more individuals, the overall detection capability of the group increases (Fig 3B), due to both having a full coverage of the surrounding area as well as having multiple overlapping visual areas for detection redundancy. However, blockage effects cause this trend to be sub-linear with respect to the number of individuals in the group (Fig 5C). This demonstrates that one of the benefits of being part of a group - the “many eyes” effect [7, 8] - has a decreasing marginal utility as group size continues to increase. To explore possible functional consequences of this, consider that individuals in a group need to not only detect an object, but also respond to the detection. While a predator may elicit a sudden startle response [20], movement towards a potential food source is more likely to be gradual. Previous work has shown that only a small fraction of ‘informed’ group members (e.g. group members that can detect the food source) are needed in order to successfully guide the group towards the target [19, 26, 38]. Here, we note that although the fraction of informed individuals needed to lead the group decreases with *N*, the average detection capability of each individual also decreases with *N*. Therefore, we can not generalize to say whether small or large groups are expected to have a better ability to both detect and move towards a promising food source, since the scaling of detection capability with *N* depends on the characteristics of individuals and the configuration of the group.

In our calculations, we considered that an individual can detect an outside point in a given direction if they have a clear view in that direction. However, this does not take into account differences in detection capability for near versus far away objects that arise due to visual projection and contrast effects. Real objects have a finite size and thus the total angle subtended by the object decreases with distance. An object located close to the group thus projects onto a larger range of angular directions compared to the same object located farther away. This naturally results in a lower overall detection capability if an object is located farther away. In addition, the contrast an object appears at with respect to the background decreases with distance due to the effects of light scattering and absorption. This can have a significant effect in attenuating media such as water, and can be particularly strong in conditions of poor visibility (e.g. in turbid, or ‘cloudy’, water - see [39, 40]). A decrease in contrast with distance could have two effects on group detection ability. First, it would lower the effective detection capability for each individual in the group. In the model, this is represented by increasing the effective baseline visual blocking probability λ_0_ (Fig 5E). Second, because visual detection only occurs if an object appears above a certain contrast threshold [40], individuals may be able to detect an object if they are close to it (i.e. located on the side of the group where the object is located), but individuals on the other side of the group may not have sufficient contrast to detect the object. Since such mechanisms represent individuals as ‘imperfect sensors’, they affect the ‘many eyes’ abilities of the group: while a group with a small number of individuals could be certain to detect an object in a condition of clear visibility, the same group may not have any individuals that detect the object in conditions of poor visibility. In a group with a larger number of individuals, the pure increase in numbers makes it is more likely at least some individuals are able to detect an object even in conditions of poor visibility. This is similar to the ‘pool-of-competence’ effect, whereby larger groups effectively act as better problem solvers because it is more likely they contain an individual that has the knowledge needed to solve the problem [41, 42]. Applying this to the case of visual detection, a larger group is more likely to contain an individual that can detect the object.

While we obtained data from freely-moving groups of fish, we note that the effective transition point from homogeneous to heterogeneous visual information available among individuals will be different for groups of different animals. Based on our results, we would expect differences due to the shape of the animal, the spacing between individuals in the group, and the overall space that the group occupies. For example, while we studied fish moving in a planar configuration in shallow water, and approximated out-of-plane effects using probabilistic visual blockage, we expect that fish moving in a fully three-dimensional (non-planar) shape would have overall a smaller fraction of their vision blocked by neighbors, for a group containing the same number of individuals. However, experiments also show that fish schools in open water often adopt planar structures, in particular in response to a nearby predator [43], and that using two-dimensional motion coordinates yields the same results for leader-follower dynamics as considering full 3-dimensional motion [44]. Other animals that form non-planar groups [45], such as midges or birds, can differ in the effective visual blockage due to neighbors. In a midge swarm, where the inter-individual spacing relative to body size is larger than that for fish [46], we would expect relatively low visual blockage. Different from fish, we would also expect minimal directional dependence, due to the body shape of midges. In contrast to midges, birds have elongated body shapes, and therefore we could expect similar direction-dependent detection trends for birds as we found for the fish schools studied here; in addition, although birds move in 3D, data from starlings has shown that flocks are generally thinnest in the direction of gravity and therefore also have planar characteristics [47]. Ungulates moving in a herd, such as zebra, gazelles, caribou, or wildebeest (e.g. [48, 49]), have both elongated body shapes and move on a two-dimensional surface, and thus may have directly comparable trends for visual detection as fish moving in shallow water.

In summary, we used fish as a model system to examine the visual information available to individuals in the group, and formulated a simple model to understand how visual information changes with number of individuals in the group. In future work it will be valuable to compare results to other animal groups that vary in their individual properties and group dynamics, and to test the expected changes in detection ability with respect to individual placement and group motion direction.

## 4 Methods

### 4.1 Experiments

Golden shiners (*Notemigonus crysoleucas*) are a small minnow native to the northeastern U.S. and Canada [27]. Juvenile shiners approximately 5 cm in length were purchased from Anderson Farms (www.andersonminnows.com) and were allowed to acclimate to the lab for two months prior to experiments. Fish were stored in seven 20-gallon tanks at a density of approximately 150 fish per tank. Tank water was conditioned, de-chlorinated, oxygenated, and filtered continuously. Fifty percent of tank water was exchanged twice per week. Nitrates, nitrites, pH, saline and ammonia levels were tested weekly. The room temperature was controlled at 16 °C, with 12 hr of light and 12 hr of dark, using dawn-dusk simulating lights. Fish were fed three times daily with crushed flake food and experiments were conducted 2-4 hr after feeding. These methodologies are identical to those used in [50].

Trials with groups of 10, 30, and 70 shiners (3 trials each) and with 151 shiners (1 trial) were allowed to swim freely in a 2.1 × 1.2m experimental tank. Water depth was 4.5 - 5 cm. Fish were filmed for 2 hr from a Sony EX-1 camera place 2m above the tank, filming at 30 frames per second.

The arena was acoustically and visually isolated from external stimuli: two layers of sound insulation were placed under the tank, and the tank was enclosed in a tent of featureless white sheets. Trials took place in a quiet laboratory with no people present during filming. All experimental procedures were approved by the Princeton University Institutional Animal Care and Use Committee.

### 4.2 Tracking and group area

We focused analysis on a 13 minute segment for each trial. We chose a time 1 hr after the onset of the trial to minimize stress on the fish from handling. Fish positions, orientations, and body postures were extracted from videos via the SchoolTracker algorithm used in [20]. Briefly, SchoolTracker works by detecting fish in each frame, then creating tracks by linking detected fish across frames. We then performed manual data correction to ensure accuracy in the tracks.

We used a convex hull and Voronoi tessalation to quantify the overall spatial area occupied by the group as well as the spatial area per individual (Fig 6).

**Figure 6:**
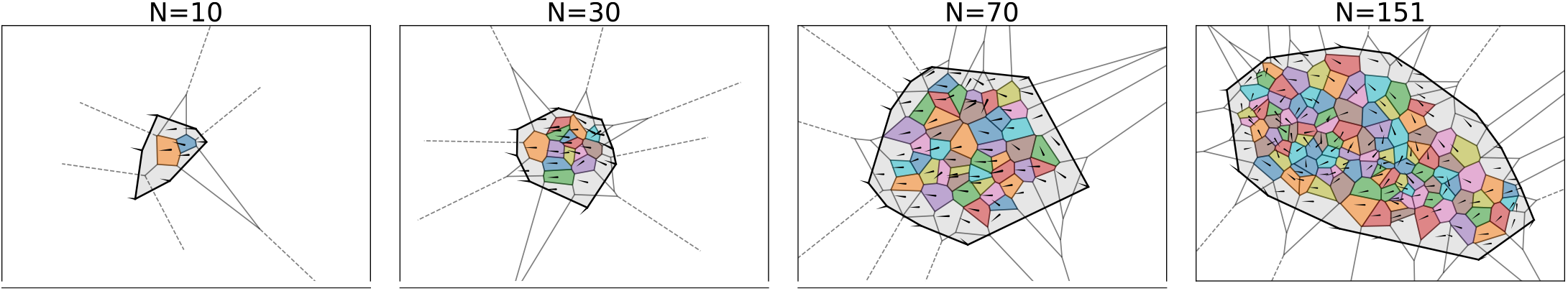
Group and individual area calculations. (A) The area of the group is calculated by a convex hull that contains the head positions of all group members (gray shading). Individual area is calculated using a Voronoi tesselation, keeping only Voronoi polygons that are enclosed in the overall group boundaries (colored areas).

### 4.3 External detection

To examine external detection, we represent individuals as simplified 4-sided polygons defined by their head, eyes, and tail (Figs 1A and 7A). To examine detection, we consider 200 points located in a circular arrangement at a distance of 1200 pixels (~ 135 cm, or ~ 30 body lengths) from the group centroid. This distance is sufficiently far from the edges of the group that the exact value does not significantly affect results. An individual has direction capability in a certain direction if there is a clear visual line in that direction, without blockage from neighbors.

To determine external detection coverage, at each time step we first shift coordinates such that the group centroid is at the origin, and follow this by a rotation that sets the direction of travel of the group centroid to be along the x-axis. In this coordinate system, each individual’s location is defined by a front-back distance *ξ_i_*(*t*) along the x-axis, and a side-side distance *ν_i_*(*t*) along the y-axis (Fig 7A). The edge of the group in each direction is defined as the individual furthest away in that direction; we denote these values as *ξ_F_* (*t*), *ξ_B_*(*t*), *ν_L_*(*t*), and *ν_R_*(*t*) for the front, back, left, and right edges, respectively (Fig 7A). In the (*ξ, ν*) coordinate system we expect side-to-side symmetry for reflections about the *ξ*-axis. However, due to both eye positions being located at the head, and a “blind angle” where individuals cannot see behind themselves, there is no front-back symmetry. We used a blind angle value of 25^*o*^, which was obtained from a visual study of our study species [29].

**Figure 7:**
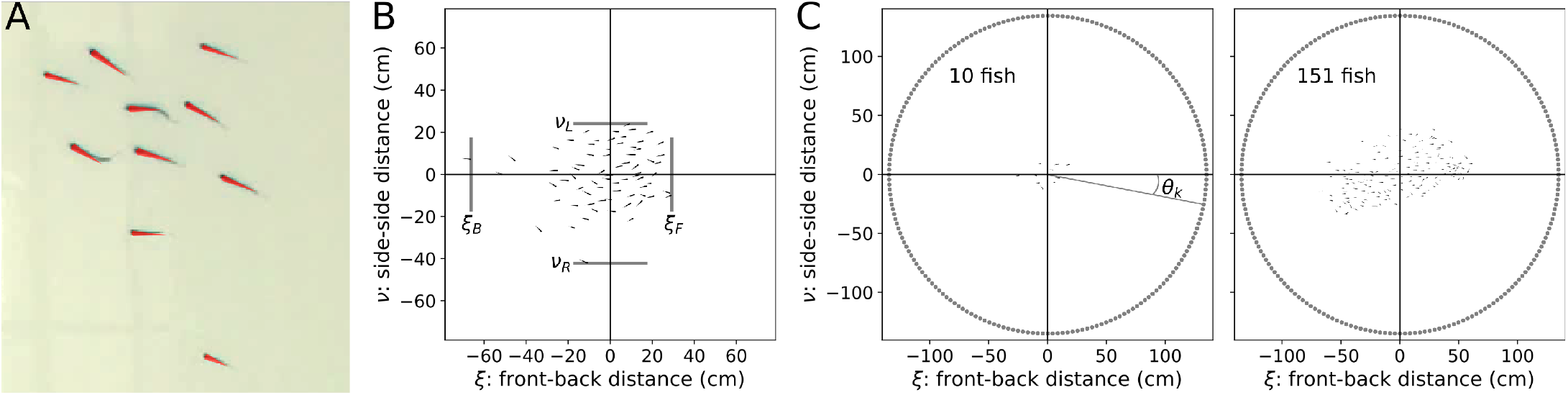
Polygon representation of fish and detection analysis quantities. (A) Example zoomed-in video frame from a group of 10 fish with the 4-sided fish polygon model shown as the red overlay. (B) By setting the origin at the group centroid and the group travel direction along the *x*-axis, we define the (*ξ, ν*) coordinate system. The front-back coordinate is *ξ*, and the side-side coordinate is *ν*. The front, back, left side, and right side of the group (*ξ_F_*, *ξ_B_*, *ν_L_*, and *ν_R_*, respectively) are defined as the head position of the individual farthest away from the group centroid in that direction. The orange arrow denotes the group direction of travel (along the x-axis) (C) To examine the angular dependence of detection, we consider 200 ‘points’ placed at a distance of 1200 pixels (~ 135 cm, or ~ 30 body lengths) from the group centroid. This distance is larger that the space occupied by the largest group. The angle *θ_k_* defines the angular location of an external point with respect to the group travel direction.

Fig 7B shows both a small and large group with the ‘circle’ of points surrounding it. Each point has an angular location *θ_k_* relative to the direction of travel of the group centroid, where *k* = 1‥200. Individual *i* has both left and right eyes located to the sides of its body, the positions of which where estimated from the tracking software. We say that individual *i* has detection capability at relative angle *θ_k_* at time *t* if there is no visual blockage between either its left eye or its right eye and the point at *θ_k_*. This defines the function

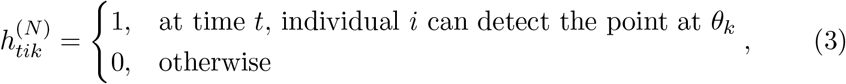

which is calculated for each group containing a different number (*N*) of individuals.

For probabilistic detection, which we use to approximate out-of-plane effects, we use an analogous calculation to that described above, but instead define a probability that the presence of a neighbor in a certain direction blocks external vision in that direction. This is done with consistent random draws that affect the left eye and right eye together. Detection is defined if an external vision path exists from either the left eye or the right eye. To consider the blind angle to the rear of of an individual, we simply exclude directions within the blind angle and mark them as not detected. Although other than the blind angle we did not place an explicit limit on the range of the left eye versus the right eye in the detection calculations, an individual’s own body blocks vision to their the opposite side, so that the left eye does not have a clear visual path to the right side, and vice versa.

We use Eq. 3 to calculate the distributions of individual detection coverage and the number of group detections. Using (·) to represent an average over the specified indices, first we define the following notation to simplify the calculations of individual detection:

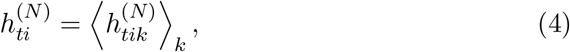

which is the individual detection at an instant in time, calculated by averaging over all possible detection directions *k*. The average individual detection coverage is

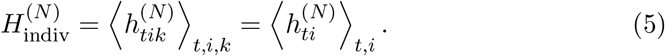

The variance of individual detection coverage is

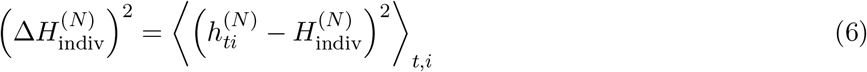

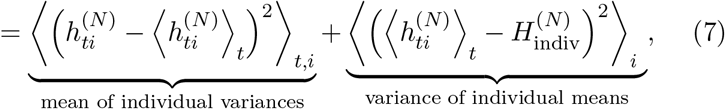

where the second line follows because the individual and temporal differences are symmetric about the mean. The first term in Eq. 7 is the mean of the individual variances, which is associated with individuals having different values of detection coverage during the course of a trial. The second term in Eq. 7 is the variance of the individual means, which is associated with consistent individual differences. Applying this to the data with results from incomplete blockage (75%) and blind angle, we obtain that the variance of the individual means explains (8.2%, 7.7%, 6.4%, 9.0%) of the total variance for the group sizes of *N* = (10, 30, 70, 151), respectively, with the remaining fraction of the variance accounted for by the mean of the individual variances.

The average number of group detections is

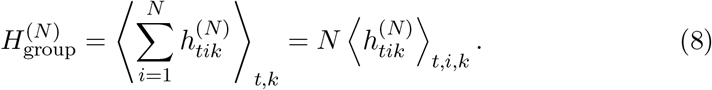

Note that while the averages in Eqs. 5 and 8 are related by the simple formula 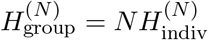, the distributions of individual and group detections do have such a simplne ortelation to each other. The standard deviation of the number of group detections is

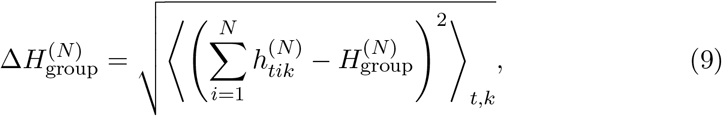

which is shown in Figs 3B and 5C.

To compute detection with respect to the group direction of travel in Fig 4, we use only polarized group states where the direction of travel is well-defined. To categorize when the group is in a polarized state or the other states shown in Fig 3, we use the same definitions as in [31].

### 4.4 Model

We formulate a simple model to describe the external visual detection coverage of individuals in a group. In this model, the group occupies a circular area with radius *R*, within which there is a constant visual blockage probability. Using symmetry, an individual’s field of view depends solely on its distance *r* from the center of group, where 0 ≤ *r* ≤ *R*. Whether or not an individual located at *r* can see outside the group in a direction *θ* depends on the distance *g*(*r, θ*) from the individual to the edge of the group in that direction (Fig 5A). Using the law of cosines, this distance is

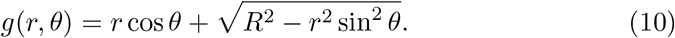

We say that the probability of being able to see outside the group in a given direction is the product of the blockage probability λ times the distance to the edge in that direction. Assuming that blocking events are randomly distributed and occur with a uniform probability through the group, we use the Poisson distribution to represent the probability of external detection:

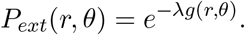

For the individual at position *r*, the total external detection capability is an average, calculated by the integral over all possible angles:

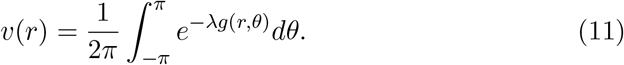

To perform calculations, we set *R* = 1, which is done without loss of generality because detection in Eq. 11 depends on the product in the exponent.

Thus far we have the assumed that the group occupies a fixed area defined by the radius *R*. However, in the data we observe that groups change the area they occupy over the course of a trial (Fig 3C). To represent a distribution of the area occupied, consider a group at two different sizes: *R* (the average radius), and *R*_1_ (the ‘current’ radius). Defining the ratio *α* = *R/R*_1_, the distance to the edge of the group scales as *g*_1_(*r*_1_*, θ*) = *g*(*r, θ*)*/α*. For the blockage probability we expect this to scale with the density within the group, and thus have λ_1_ = *λα*^2^. For a current state of the group defined by the size ratio *α*, the external visual detection coverage of an individual is calculated by using Eq. 11 in the current state:

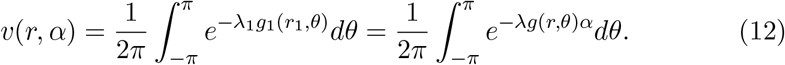

For simplicity, we represent different group areas by using a Gaussian with a mean of *α* = 1 to represent different possible values of the group radius,

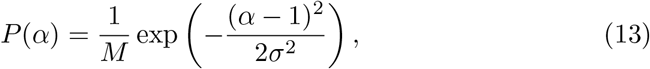

where *σ* represents the magnitude of changes in the group radius, and *M* is a normalization factor. Because the radius must be positive, we restrict to values *α* > 0.

To compute the probability distribution of external detection, we evaluate Eq. 12 on a discrete set of radii, calculating the number of individuals in a shell around a given value of *r* as proportional to

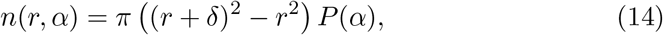

where *δ* is the width of the shell. We then use binning to calculate the probability distribution of detection coverage, using Eq. 13 to represent the probability of different group areas. To apply the model to the groups with different numbers of individuals, we specify that λ varies to a power of the number of individuals (*N*) in the group (main text Eq. 1, repeated here):

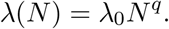

The model results for all values of *N* are defined by the three parameters λ_0_, *q*, and *σ*. Because the average detection coverage depends only very weakly on the value of *σ*, we use a two-step procedure to fit these parameters to the data, fitting λ_0_ and *q* to the average detection coverage, and then subsequently fitting *σ* to the standard deviation of the number of group detections.

Note that individuals maintaining constant density with an increase in *N* can be approximated with *q* = 0.5 (this represents a linear increase in area with *N*). However, even if individuals did maintain constant density, this scaling would only be strictly true for point particles. To see why, consider the case where individuals are zero-dimensional ‘points’; then, the visual blockage probability would only depend on the density of points, and would be constant with distance for uniformly distributed points. However, since instead a group member has a 2-dimensional projection represented in our calculations by a polygon, the visual blockage probability depends both on the density of neighbors and the distance from each observer. Because of this, we fit both the values of λ_0_ and *q*. The fitting procedure for these parameters minimizes the mean square error of the model result for mean external detection capability compared to the data for each value of *N*, where values from the data are used from the incomplete blocking detection procedure (Fig 5B).

Following this, we fit *σ* by minimizing the mean square error of the model result for the standard deviation of the number of group detections compared to the data (Fig 5C). Note that in the model, a single “snapshot” of the group is defined by a particular value of the group area.

#### 4.4.1 Analytical approximation for average detection capability

To obtain an analytical approximation, consider Eq. 11, which is the detection capability of an individual located at position *r*. Using a single value for the group area, the average visual degree is an integral over the unit sphere of Eq. 11 times the probability that an individual is located at *r*:

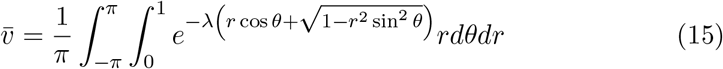

Although this cannot be evaluated in closed form, we can obtain an approximation by considering the series expansion in powers of *r*:

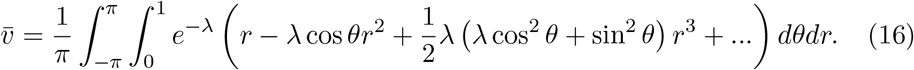

Evaluating the integral for the individual terms yields an expression in the form of an exponential times a series expansion in powers of λ (main text Eq. 2, repeated here):

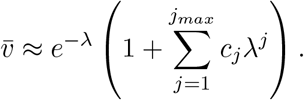

Keeping terms to 6^th^ order in *r* has *j_max_* = 6 and the following coefficient values: *c*_1_ = 0.1455, *c*_2_ = 0.1455, *c*_3_ = 1.302 × 10^−2^, *c*_4_ = 6.185 × 10^−3^, *c*_5_ = 3.255 × 10^−4^, *c*_6_ = 1.085 × 10^−4^. We use the above expression (Eq. 2) with these coefficient values to plot the series approximation in Fig 5D-E.

## 5 Acknowledgments

We thank the Couzin Laboratory for helpful discussions. In particular we would like to thank Yael Katz for help with experimental work.

## 6 Funding

We acknowledge funding from the Max Planck Institute of Animal Behavior. I.C. acknowledges support from the NSF (IOS-1355061), the Office of Naval Research (ONR, N00014-19-1-2556), the Struktur-und Innovationsfonds für die Forschung of the State of Baden-Württemberg, the Deutsche Forschungsgemeinschaft (DFG, German Research Foundation) under Germany’s Excellence Strategy– “Centre for the Advanced Study of Collective Behaviour” EXC 2117-422037984, and the Max Planck Society. J.D. acknowledges support from the Heidelberg Academy of Science, Baden-Württemberg, Germany, under the WIN (wissenschaftlichen Nachwuchs) program. M.S. acknowledges funding from the NSF Graduate Research Fellowship. C.T. was supported by a MindCORE (Center for Outreach, Research, and Education) Postdoctoral Fellowship.

## Notes

https://github.com/jacobdavidson/collectivedetection

### Competing Interest Statement

The authors have declared no competing interest.

### Summary of Updates

Added ORCID for V.H. Sridhar and I.D. Couzin.

## References

1. WD Hamilton. Geometry for the selfish herd. Journal of Theoretical Biology, 31(2):295–311, 1971.

2. Benoni H. Seghers. Schooling Behavior in the Guppy (Poecilia reticulata): An Evolutionary Response to Predation. Evolution, 28(3):486–489, 1974. Publisher: [Society for the Study of Evolution, Wiley].

3. Tony J. Pitcher. Functions of Shoaling Behaviour in Teleosts. In Tony J. Pitcher, editor, The Behaviour of Teleost Fishes, pages 294–337. Springer US, Boston, MA, 1986.

4. Anne E. Magurran. The adaptive significance of schooling as an anti-predator defence in fish. Annales Zoologici Fennici, 27(2):51–66, 1990. Publisher: Finnish Zoological and Botanical Publishing Board.

5. A Berdahl, CJ Torney, CC Ioannou, JJ Faria, and ID Couzin. Emergent sensing of complex environments by mobile animal groups. Science, 339(6119):574–576, 2013.

6. Andrew M. Hein, Sara Brin Rosenthal, George I. Hagstrom, Andrew Berdahl, Colin J. Torney, and Iain D. Couzin. The evolution of distributed sensing and collective computation in animal populations. eLife, 4:e10955, December 2015.

7. SL Lima. Back to the basics of anti-predatory vigilance: the group-size effect. Animal Behaviour, 49:11–20, 1995.

8. Steven L. Lima. The influence of models on the interpretation of vigilance. Readings in animal cognition. MIT Press, Cambridge, Massachusetts, USA, page 201–216, 1996.

9. Guy Beauchamp. Reduced flocking by birds on islands with relaxed predation. Proceedings of the Royal Society of London. Series B: Biological Sciences, 271(1543):1039–1042, May 2004.

10. Nikolai W. F. Bode, Jolyon J. Faria, Daniel W. Franks, Jens Krause, and A. Jamie Wood. How perceived threat increases synchronization in collectively moving animal groups. Proceedings of the Royal Society B: Biological Sciences, 277(1697):3065–3070, October 2010.

11. DJ Hoare, ID Couzin, J-GJ Godin, and J Krause. Context-dependent group size choice in fish. Animal Behaviour, 67(1):155–164, 2004.

12. Mary C. Hager and Gene S. Helfman. Safety in numbers: shoal size choice by minnows under predatory threat. Behavioral Ecology and Sociobiology, 29(4):271–276, November 1991.

13. Marko Spieler and Karl Eduard Linsenmair. Aggregation Behaviour of Bufo maculatus Tadpoles as an Antipredator Mechanism. Ethology, 105(8):665–686, 1999.

14. Matthew M. G. Sosna, Colin R. Twomey, Joseph Bak-Coleman, Winnie Poel, Bryan C. Daniels, Pawel Romanczuk, and Iain D. Couzin. Individual and collective encoding of risk in animal groups. Proceedings of the National Academy of Sciences, 116(41):20556–20561, October 2019.

15. S Creel, P Schuette, and D Christianson. Effects of predation risk on group size, vigilance, and foraging behavior in an african ungulate community. Behavioral Ecology, 25:773–784, 2014.

16. S Creel and WA Winnie. Jr. Responses of elk herd size to fine-scale spatial and temporal variation in risk of predation by wolves. Animal Behaviour, 69:1181–1189, 2005.

17. JE Orpwood, AE Magurran, JD Armstrong, and SW Griffiths. Minnows and the selfish herd: effects of predation risk on shoaling behaviour are dependent on habitat complexity. Animal Behaviour, 76:143–152, 2008.

18. Lesley J. Morrell, Graeme D. Ruxton, and Richard James. Spatial positioning in the selfish herd. Behavioral Ecology, 22(1):16–22, January 2011.

19. A Strandburg-Peshkin, CR Twomey, NWF Bode, AB Kao, Y Katz, CC Ioannou, SB Rosenthal, CJ Torney, HS Wu, SA Levin, and ID Couzin. Visual sensory networks and effective information transfer in animal groups. Current Biology, 23(17):R709–R711, 2013.

20. SB Rosenthal, CR Twomey, AT Hartnett, HS Wu, and ID Couzin. Revealing the hidden networks of interaction in mobile animal groups allows prediction of complex behavioral contagion. Proceedings of the National Academy of Sciences, 112(15):4690–4695, 2015.

21. J Parrish and L Edelstein-Keshet. Complexity, pattern, and evolutionary trade-offs in animal aggregation. Science, 284(2):99–101, 1999.

22. J Krause and GD Ruxton. Living in groups. Oxford University Press, Oxford, 2002.

23. K. Kotrschal, M.J. Van Staaden, and R. Huber. Fish Brains: Evolution and Anvironmental Relationships. Reviews in Fish Biology and Fisheries, 8(4):373–408, December 1998.

24. Martin Egelhaaf and Roland Kern. Vision in flying insects. Current Opinion in Neurobiology, 12(6):699–706, December 2002.

25. Michael P. Jones, Kenneth E. Pierce, and Daniel Ward. Avian Vision: A Review of Form and Function with Special Consideration to Birds of Prey. Journal of Exotic Pet Medicine, 16(2):69–87, April 2007.

26. Iain D. Couzin, Christos C. Ioannou, Güven Demirel, Thilo Gross, Colin J. Torney, Andrew Hartnett, Larissa Conradt, Simon A. Levin, and Naomi E. Leonard. Uninformed individuals promote democratic consensus in animal groups. Science, 334(6062):1578–1580, 2011.

27. TR Whittier, DB Halliwell, and RA Daniels. Distributions of lake fishes in the northeast - ii. the minnows (cyprinidae). Northeastern Naturalist, 7(2):131–156, 2000.

28. Nathan M. Stone, Anita M. Kelly, and Luke A. Roy. A Fish of Weedy Waters: Golden Shiner Biology and Culture. Journal of the World Aquaculture Society, 47(2):152–200, 2016.

29. Diana Pita, Bret A. Moore, Luke P. Tyrrell, and Esteban Fernández-Juricic. Vision in two cyprinid fish: implications for collective behavior. PeerJ, 3:e1113, August 2015.

30. Simon Leblanc. Information Flow on Interaction Networks. PhD thesis, Princeton University, 2018.

31. K Tunstrøm, Y Katz, CC Ioannou, C Huepe, MJ Lutz, and ID Couzin. Collective states, multistability and transitional behavior in schooling fish. PLoS Computational Biology, 9(2), 2013.

32. J Krause. Differential fitness returns in relation to spatial position in groups. Biological Reviews, 69:187–206, 1994.

33. Damon Centola. The Spread of Behavior in an Online Social Network Experiment. Science, 329(5996):1194–1197, September 2010. Publisher: American Association for the Advancement of Science Section: Report.

34. Daniel M. Romero, Brendan Meeder, and Jon Kleinberg. Differences in the mechanics of information diffusion across topics: idioms, political hashtags, and complex contagion on twitter. In Proceedings of the 20th international conference on World wide web, WWW’11, pages 695–704, New York, NY, USA, March 2011. Association for Computing Machinery.

35. Damon Centola. How Behavior Spreads: The Science of Complex Contagions. Princeton University Press, Princeton, NJ, March 2020. Google-Books-ID: 6SCyDwAAQBAJ.

36. Josh A. Firth. Considering Complexity: Animal Social Networks and Behavioural Contagions. Trends in Ecology – Evolution, 35(2):100–104, February 2020.

37. Hannah E. A. MacGregor, James E. Herbert-Read, and Christos C. Ioannou. Information can explain the dynamics of group order in animal collective behaviour. Nature Communications, 11(1):2737, June 2020. Number: 1 Publisher: Nature Publishing Group.

38. Iain D. Couzin, Jens Krause, Nigel R. Franks, and Simon A. Levin. Effective leadership and decision-making in animal groups on the move. Nature, 433(7025):513–516, February 2005.

39. Thomas W. Cronin, Sönke Johnsen, N. Justin Marshall, and Eric J. Warrant. Visual Ecology. Princeton University Press, August 2014. Publication Title: Visual Ecology.

40. Colin Robert Twomey. Vision and motion in collective behavior. PhD thesis, Princeton University, 2016.

41. Julie Morand-Ferron and John L. Quinn. Larger groups of passerines are more efficient problem solvers in the wild. Proceedings of the National Academy of Sciences, 108(38):15898–15903, September 2011.

42. Max Wolf and Jens Krause. Why personality differences matter for social functioning and social structure. Trends in Ecology – Evolution, 29(6):306–308, June 2014.

43. Maksym Romenskyy, James E Herbert-Read, Christos C Ioannou, Alex Szorkovszky, Ashley J W Ward, and David J T Sumpter. Quantifying the structure and dynamics of fish shoals under predation threat in three dimensions. Behavioral Ecology, 31(2):311–321, March 2020.

44. Isobel Watts, Máté Nagy, Robert I. Holbrook, Dora Biro, and Theresa Burt de Perera. Validating two-dimensional leadership models on three-dimensionally structured fish schools. Royal Society Open Science, 4(1):160804, January 2017.

45. Julia K. Parrish and William M. Hamner. Animal Groups in Three Dimensions: How Species Aggregate. Cambridge University Press, Dec 1997.

46. Douglas H. Kelley and Nicholas T. Ouellette. Emergent dynamics of laboratory insect swarms. Scientific Reports, 3(1):1–7, Jan 2013.

47. Michele Ballerini, Nicola Cabibbo, Raphael Candelier, Andrea Cavagna, Evaristo Cisbani, Irene Giardina, Alberto Orlandi, Giorgio Parisi, Andrea Procaccini, Massimiliano Viale, and Vladimir Zdravkovic. Empirical investigation of starling flocks: a benchmark study in collective animal behaviour. Animal Behaviour, 76(1):201–215, July 2008.

48. Shay Gueron and Simon A. Levin. Self-organization of front patterns in large wildebeest herds. Journal of theoretical Biology, 165(4):541–552, 1993.

49. Torney Colin J., Lamont Myles, Debell Leon, Angohiatok Ryan J., Leclerc Lisa-Marie, and Berdahl Andrew M. Inferring the rules of social interaction in migrating caribou. Philosophical Transactions of the Royal Society B: Biological Sciences, 373(1746):20170385, May 2018.

50. Y Katz, K Tunstrøm, CC Ioannou, C Huepe, and ID Couzin. Inferring the structure and dynamics of interactions in schooling fish. Proceedings of the National Academy of Sciences, 108:18720–18725, 2011.

